# Non-ionotropic NMDA receptor signaling gates bidirectional structural plasticity of dendritic spines

**DOI:** 10.1101/2020.08.19.258269

**Authors:** Ivar S. Stein, Deborah K. Park, Nicole Claiborne, Karen Zito

## Abstract

Experience-dependent refinement of neuronal connections is critically important for brain development and learning. Here we show that ion flow-independent NMDAR signaling is required for the long-term dendritic spine growth that is a vital component of brain circuit plasticity. We found that inhibition of p38 MAPK, shown to be downstream of non-ionotropic NMDAR signaling in LTD and spine shrinkage, blocked LTP-induced spine growth but not LTP. We hypothesized that non-ionotropic NMDAR signaling drives the cytoskeletal changes that support bidirectional spine structural plasticity. Indeed, we found that key signaling components downstream of non-ionotropic NMDAR function in LTD-induced spine shrinkage also are necessary for LTP-induced spine growth. Furthermore, NMDAR conformational signaling with coincident Ca^2+^ influx is sufficient to drive CaMKII-dependent long-term spine growth, even when Ca^2+^ is artificially driven through voltage-gated Ca^2+^ channels. Our results support a model in which non-ionotropic NMDAR signaling gates the bidirectional spine structural changes vital for brain plasticity.

## INTRODUCTION

Dynamic alternations in neuronal connectivity are critical for the experience-dependent modification of brain circuits throughout development and during learning. In particular, the bidirectional structural plasticity of dendritic spines is a vital process in the refinement of synaptic circuits in the mammalian cortex (e.g. (Hayashi-Takagi et al., 2015; Lai et al., 2018). Increases in synaptic strength, as through the induction of long-term potentiation (LTP), are associated with spine enlargement and new spine formation (Matsuzaki et al., 2004; Nishiyama and Yasuda, 2015), while decreases in synaptic strength, as through the induction of long-term depression (LTD), are associated with spine shrinkage or loss (Zhou et al., 2004; Stein and Zito, 2019). Notably, activation of the NMDA-type glutamate receptor (NMDAR) is required for both the spine growth associated with LTP and the spine shrinkage associated with LTD.

Recent studies have demonstrated that LTD (Nabavi et al., 2013; Carter and Jahr, 2016; Wong and Gray, 2018) and dendritic spine shrinkage (Birnbaum et al., 2015; Stein et al., 2015; Thomazeau et al., 2020) can occur independent of ion flux through the NMDA receptor. These findings have supported a model in which glutamate binding leads to conformational changes in the NMDAR that drive spine shrinkage and synaptic weakening. Indeed, imaging studies using FRET reporters have shown that NMDA or glutamate binding triggers conformational changes in the NMDAR intracellular domains, and changes its interaction with calcium/calmodulin-dependent protein kinase II (CaMKII) and protein phosphatase 1 (PP1) (Aow et al., 2015; Dore et al., 2015; Ferreira et al., 2017). p38 MAPK has been identified as a key component of the molecular pathway downstream of NMDAR conformational signaling (Nabavi et al., 2013; Stein et al., 2015), and a recent study further identified nNOS, nNOS-NOS1AP interactions, MAPK-activated protein kinase 2 (MK2) and cofilin as part of this signaling pathway (Stein et al., 2020).

Here, we made the surprising discovery that ion flow-independent NMDAR signaling is required for the long-term spine growth associated with synaptic strengthening. We show that p38 MAPK, an essential signaling component for LTD and spine shrinkage, is also required for spine growth, but not LTP, suggesting that non-ionotropic NMDAR signaling is a vital component of bidirectional spine structural plasticity. Indeed, we further show that key components of conformational NMDAR signaling, including interaction between NOS1AP and nNOS, MK2, nNOS, and CaMKII activity are all required for LTP-induced spine growth. Importantly, we also demonstrate that, when combined with non-ionotropic NMDAR signaling, long-term spine growth can be driven by Ca^2+^ influx through voltage-gated Ca^2+^ channels. Our findings support a model whereby non-ionotropic NMDAR signaling leads to disruption of the spine F-actin network, which drives spine shrinkage unless coincident Ca^2+^-influx converts Factin remodeling instead to growth.

## MATERIALS AND METHODS

All experimental protocols were approved by the University of California Davis Institutional Animal Care and Use Committee.

### Preparation and transfection of organotypic hippocampal slice cultures

Organotypic hippocampal slices were prepared from P6-P8 Sprague-Dawley rats or C57BL/6J mice of both sexes, as described (Stoppini et al., 1991). Cultures were transfected 1-2 d (EGFP) or 2 d (GCaMP6f) before imaging via biolistic gene transfer (180 psi), as previously described (Woods and Zito, 2008). 10-15 μg of EGFP-n1 or 5 μg GCaMP6f (gift from Lin Tian and Karel Svoboda; (Chen et al., 2013)) and 10 μg pCAG-CyRFP1 (Laviv et al., 2016) were coated onto 6-8 mg of 1.6 μm gold beads.

### Preparation of acute hippocampal slices

Acute hippocampal slices were prepared from P16-P20 GFP-M mice (Feng et al., 2000) of both sexes. Coronal 400 μm slices were cut (Leica VT100S vibratome) in cold choline chloride dissection solution containing (in mM): 110 choline chloride, 2.5 KCl, 25 NaHCO_3_, 0.5 CaCl_2_, 7 MgCl_2_, 1.3 NaH_2_PO_4_, 11.6 sodium ascorbate, 3.1 sodium pyruvate, and 25 glucose, saturated with 95% O_2_/5% CO_2_. Slices were recovered first at 30°C for 45 min and then at room temperature for an additional 45 min, in oxygenated artificial cerebrospinal fluid (ACSF) containing (in mM): 127 NaCl, 25 NaHCO_3_, 1.25 NaH_2_PO_4_, 2.5 KCl, 25 glucose, 2 CaCl_2_, and 1 MgCl_2_, before imaging experiments were initiated.

### Time-lapse two-photon imaging

Transfected CA1 pyramidal neurons [14-18 days in vitro (DIV)] at depths of 10-50 μm were imaged using a custom two-photon microscope (Woods et al., 2011) controlled with ScanImage (Pologruto et al., 2003). Image stacks (512 × 512 pixels; 0.02 μm per pixel) with 1-μm z-steps were collected. For each neuron, one segment of secondary or tertiary basal dendrite was imaged at 5 min intervals at 29 °C in recirculating artificial cerebral spinal fluid (ACSF; in mM: 127 NaCl, 25 NaHCO_3_, 1.2 NaH_2_PO_4_, 2.5 KCl, 25 D-glucose, aerated with 95%O_2_/5%CO_2_, ~310 mOsm, pH 7.2) with 1 μM TTX, 0 mM Mg^2+^, and 2 mM Ca^2+^, unless otherwise stated. 10 μM L-689,560 (L-689, 15 mM stock in DMSO), 100 μM 7CK (100 mM stock in H_2_O), 2 μM SB203580 (4 mM stock in DMSO), 100 μM NG-Nitro-L-arginine (L-NNA, 200 mM stock in 0.25 N HCL), 10 μM Bay-K (10 mM stock in DMSO), 50 μM (RS)-CPP (50 mM stock in H_2_O), 50 μM NBQX (10 mM stock in H_2_O); 10 μM MK2 inhibitor III (20 mM stock in DMSO); and 5 μM TAT-CN21 (5 mM stock in H_2_O; (Vest et al., 2007)) and 5 μM TAT-SCR (5 mM stock in H_2_O) were included, as indicated. Slices were pre-incubated for at least 30 min with the drug or vehicle control before glutamate uncaging. Peptides were obtained from GenicBio: L-TAT-GESV: NH2-GRKKRRQRRRYAGQWGESV-COOH, L-TAT-GASA: NH2-GRKKRRQRRRYAGQWGASA-COOH. Slices were pre-incubated with 1 μM (2 mM stock in H_2_O) peptide for at least 60 min before stimulation.

### Photolysis of MNI-caged glutamate with HFU and HFU+ stimulation

High-frequency uncaging (HFU) consisted of 60 pulses (720 nm; 2 ms duration, ~7-10 mW at the sample; adjusted to evoke an average response of ~10 pA at the soma) at 2 Hz, delivered in ACSF containing (in mM): 2 Ca^2+^, 0 Mg^2+^, 2.5 MNI-glutamate, and 0.001 TTX. The beam was parked at a point ~0.5-1 μm from the spine at the position farthest from the dendrite. HFU+ stimulation was designed to increase Ca^2+^-influx through voltage-gated calcium channels. HFU+ consisted of 60 pulses (720 nm; 8 ms duration, ~7 mW at the sample) at 6 Hz, delivered in ACSF containing (in mM): 10 Ca^2+^, 0 Mg^2+^, 5 MNI-glutamate, and 0.001 TTX. Healthy and stimulus responsive cells were selected based on a test HFU stimulus, applied before the application of pharmacological reagents. Only cells with a dendritic spine that displayed transient growth in response to the test HFU were used for experiments. Spines used for experimental data collection were on a different dendrite than the test spine.

### Electrophysiology

Whole-cell recordings (V_hold_ = −65 mV; series resistance 20-40 MΩ) were obtained from visually identified CA1 pyramidal neurons in slice culture (14-18 DIV, depths of 10-50 μm) at 25°C in ACSF containing in mM: 2 CaCl_2_, 1 MgCl_2_, 0.001 TTX, 2.5 MNI-glutamate. 2 μM SB203580 was included, as indicated. Recording pipettes (~5-7 MΩ) were filled with cesium-based internal solution (in mM: 135 Cs-methanesulfonate, 10 Hepes, 10 Na2 phosphocreatine, 4 MgCl_2_, 4 Na2-ATP, 0.4 Na-GTP, 3 Na L-ascorbate, 0.2 Alexa 488, and ~300 mOsm, ~pH 7.25). For each cell, baseline uEPSCs were recorded from two spines (2-12 μm apart) on secondary or tertiary basal branches (50-120 μm from the soma). The high-frequency glutamate uncaging stimulus (720 nm, 1 ms duration, 8-10 mW at the sample) was then applied to one spine, during which the cell was depolarized to 0 mV. Following the HFU stimulus, uEPSCs were recorded from both the target and neighboring spine at 5 min intervals for 25 min.

### Calcium imaging

CA1 pyramidal neurons (13-18 DIV) in slice culture co-expressing GCaMP6 and CyRFP1 were imaged in line-scan mode (500 Hz) to assess if they were healthy and responsive using a test stimulation of a single glutamate uncaging pulse at a dendritic spine. Using a different dendritic segment than the test spine, responsive CA1 neurons were then imaged in frame-scan mode (64 pixels per line, 7.8 Hz) before and after glutamate uncaging (720 nm, 60 pulses, 8 ms duration at 6 Hz, ~7 mW at sample) adjacent to the spine head at 27 °C in ACSF containing the following (in mM): 10 Ca^2+^, 0 Mg^2+^, 5 MNI-glutamate, and 0.001 TTX. Neurons with high baseline GCaMP6f and neurons that exhibited large calcium transients extending across the dendritic shaft were excluded.

### Quantification of data from imaging experiments

Data analysis was performed blind to the experimental condition. Cells for each condition were obtained from at least three independent hippocampal acute slices or slice culture preparations. For spine structural plasticity experiments, stimulated spine volume was estimated from background-subtracted green fluorescence using the integrated pixel intensity of a boxed region surrounding the spine head, as previously described (Woods et al., 2011). Previous studies comparing the results from this method to those obtained from electron microscopy have shown that the use of integrated fluorescence intensity to be a valid method for estimating spine volume (Holtmaat et al., 2005). For calcium imaging experiments, Ca^2+^ transient peak amplitude (ΔF/F_0_) was measured from background-subtracted green fluorescence in the spine as the ratio of fluorescence during HFU (of specified windows after HFU) to basal fluorescence (2.4 s window before uncaging). All images are maximum projections of three-dimensional (3D) image stacks after applying a median filter (3 × 3) to the raw image data.

### Quantification of data from electrophysiology experiments

Data analysis was performed blind to the experimental condition. Cells for each condition were obtained from at least three independent hippocampal acute slices or slice culture preparations. uEPSC amplitudes from individual spines were quantified as the average of 5-6 test pulses at 0.1 Hz) from a 2 ms window centered on the maximum current amplitude within 50 ms following uncaging pulse delivery relative to the baseline.

### Statistical analysis

All data are represented as mean ± standard error of the mean (SEM). All statistics were calculated across cells. Statistical significance was set at p < 0.05 (two-tailed t test). Fig 1–4 utilizes two-tailed unpaired t test for the line graphs and two-tailed paired t test for the bar graphs; Fig S1C utilizes two-tailed unpaired t test and Fig S1D utilizes two-tailed paired t test. All p and n values are presented in the Results section and figure legends.

**Figure 1.**
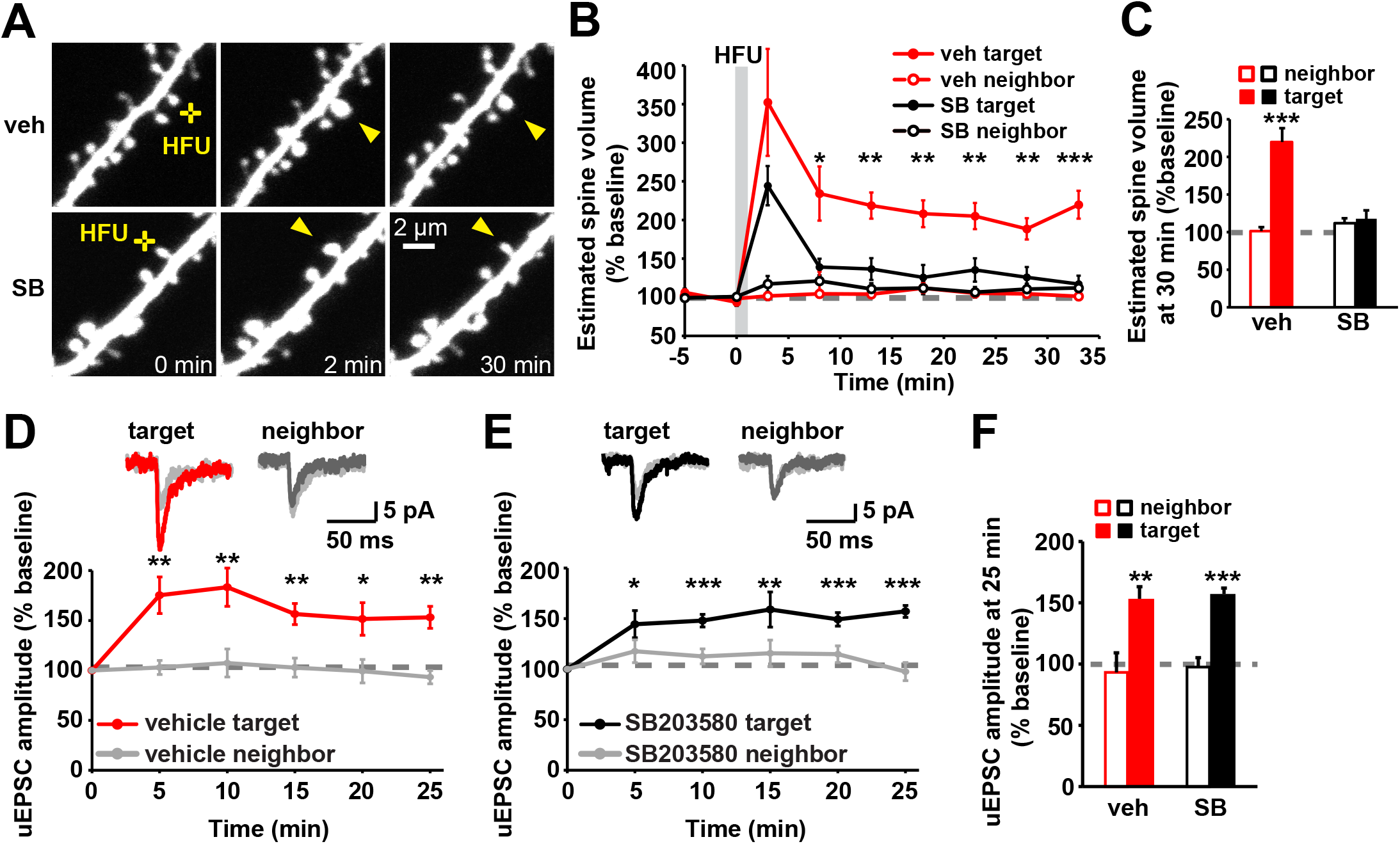
p38 MAPK activity is required for LTP-induced spine growth, but not for synaptic strengthening. **(A)** Images of dendrites from DIV14-18 EGFP-expressing CA1 neurons before and after HFU stimulation (yellow cross) at individual dendritic spines (yellow arrowhead) with and without p38 MAPK inhibitor SB203580 (SB, 2 μM). **(B, C)** Inhibition of p38 MAPK with SB (black filled circles/bar; 11 spines/11 cells) reduced HFU-induced spine growth compared to vehicle (red filled circles/bar; 11 spines/11 cells). Volume of unstimulated neighbors (open circles/bars) was unaffected. **(D, E)** Top, average traces of uEPSCs from a target spine and an unstimulated neighbor before (gray) and 25 min after HFU stimulation during vehicle conditions (target, red; neighbor, dark gray) or in the presence of SB (target, black; neighbor, dark gray). Bottom, HFU-induced increases in uEPSC amplitude in vehicle (red; 8 spines/8 cells) were unaffected by p38 MAPK inhibition with SB (black; 9 spines/9 cells). uEPSC amplitudes of unstimulated neighboring spines (gray) were unaffected. **(F)** HFU induced a long-lasting uEPSC amplitude increase (red filled bar) compared to baseline, which is unaffected by p38 MAPK inhibition (black filled bar). Data are represented as mean ± SEM. *p < 0.05; **p < 0.01, ***p < 0.001.

**Figure 2.**
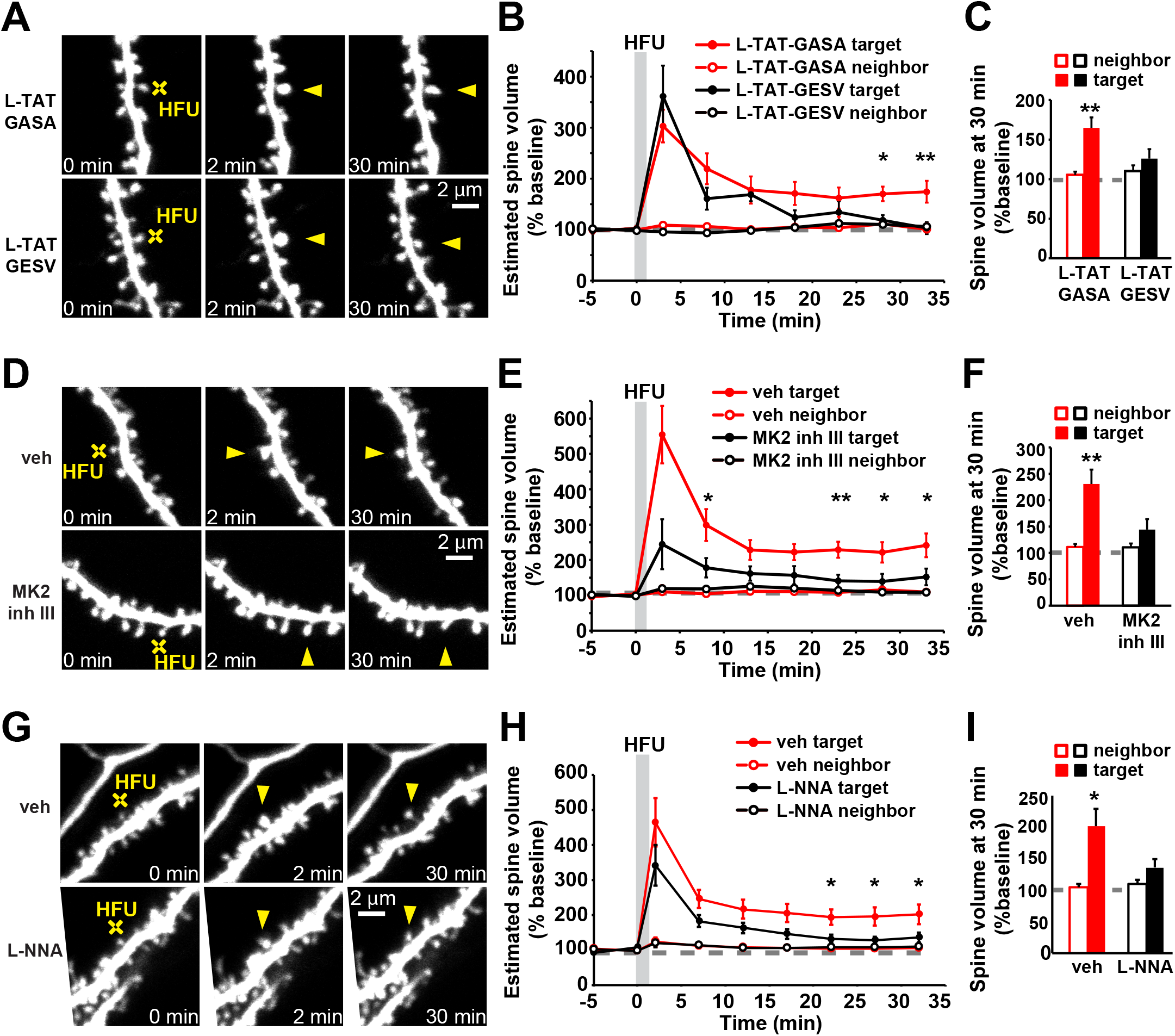
Non-ionotropic NMDAR signaling pathway is required for LTP-induced spine growth. **(A, D, G)** Images of dendrites from EGFP-expressing CA1 neurons at DIV14-18 before and after HFU stimulation (yellow cross) of individual spines (yellow arrowhead) in the presence of L-TAT-GESV (1 μM) and the L-TAT-GASA control peptide (1 μM), during vehicle conditions and in the presence of MK2 inhibitor III (10 μM) or the NOS inhibitor, L-NNA (100 μM). **(B, C)** Disruption of the NOS1AP/nNOS interaction with L-TAT-GESV (black filled circles/bar; 9 spines/9 cells), but not application of the inactive L-TAT-GASA control peptide (red filled circles/bar; 9 spines/9 cells) inhibited persistent spine growth following LTP induction. Volume of the unstimulated neighbors (open circles/bars) was unchanged. **(E, F)** Inhibition of MK2 activity (black filled circles/bar; 11 spines/11 cells) prevented HFU-induced persistent spine enlargement (red filled circles/bar; 12 spines/12 cells). Volume of the unstimulated neighbors did not change (open circles/bars). **(H, I)** Inhibition of NOS activity (black filled circles/bar; 11 spines/11 cells) prevented HFU-induced persistent spine enlargement (red filled circles/bar; 12 spines/12 cells). Volume of the unstimulated neighbors did not change (open circles/bars). Data are represented as mean ± SEM. *p < 0.05; **p < 0.01, ***p < 0.001.

**Figure 3.**
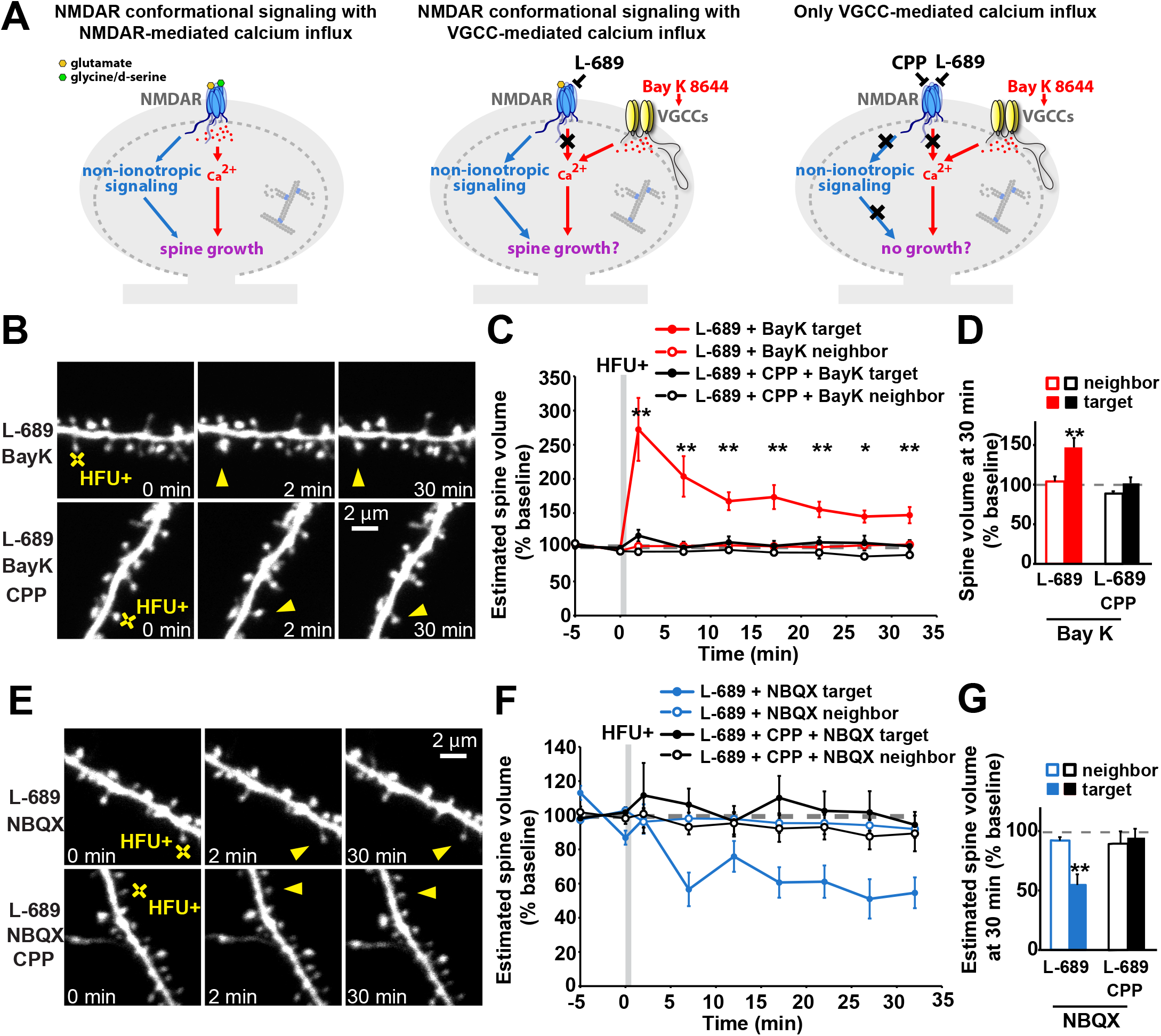
Non-ionotropic NMDAR signaling with Ca^2+^-influx through voltage-gated calcium channels is sufficient to drive LTP-induced spine growth. **(A)** Left: Proposed model whereby activity-induced spine growth requires both non-ionotropic and ionotropic NMDAR signaling. Middle: Schematic of experiment to test proposed model. Non-ionotropic NMDAR signaling is activated with glutamate while calcium influx through the NMDAR is blocked with L-689. Instead, calcium influx is driven through VGCCs with the stronger HFU+ conditions in the presence of Bay K to favor opening of VGCCs. Right: Control experiments block non-ionotropic NMDAR signaling by blocking glutamate binding to the NMDAR with CPP. **(B)** Images of dendrites from CA1 neurons of acute slices from P16-20 GFP-M mice before and after HFU+ stimulation (yellow cross) of individual spines (yellow arrowhead) in the presence of L-689 (10 μM) and Bay K (10 μM) or in combination with CPP (50 μM). **(C, D)** HFU stimulation drives spine growth in the presence of Bay K, even when ion flow through the NMDAR is blocked with L-689 (red filled circles/bar; 9 spines/9 cells), but not when non-ionotropic NMDAR signaling is blocked with CPP (black filled circles/bar; 10 spines/10 cells). Volume of the unstimulated neighbors (open circles/bars) was unchanged. **(E)** Images of dendrites under experimental conditions in B-D, except blocking the influx of calcium through VGCCs by removing Bay K and adding NBQX (50 μM). **(F, G)** Non-ionotropic NMDAR signaling led to spine shrinkage in the presence of L-689 and NBQX blue filled circles/bar; 7 spines/7 cells) but was blocked in the presence of CPP (black filled circles/bar; 6 spines/6 cells). Volume of the unstimulated neighbors (open circles/bars) did not change. Data are represented as mean ± SEM. *p < 0.05; **p < 0.01, ***p < 0.001. See also Figure S1.

**Figure 4.**
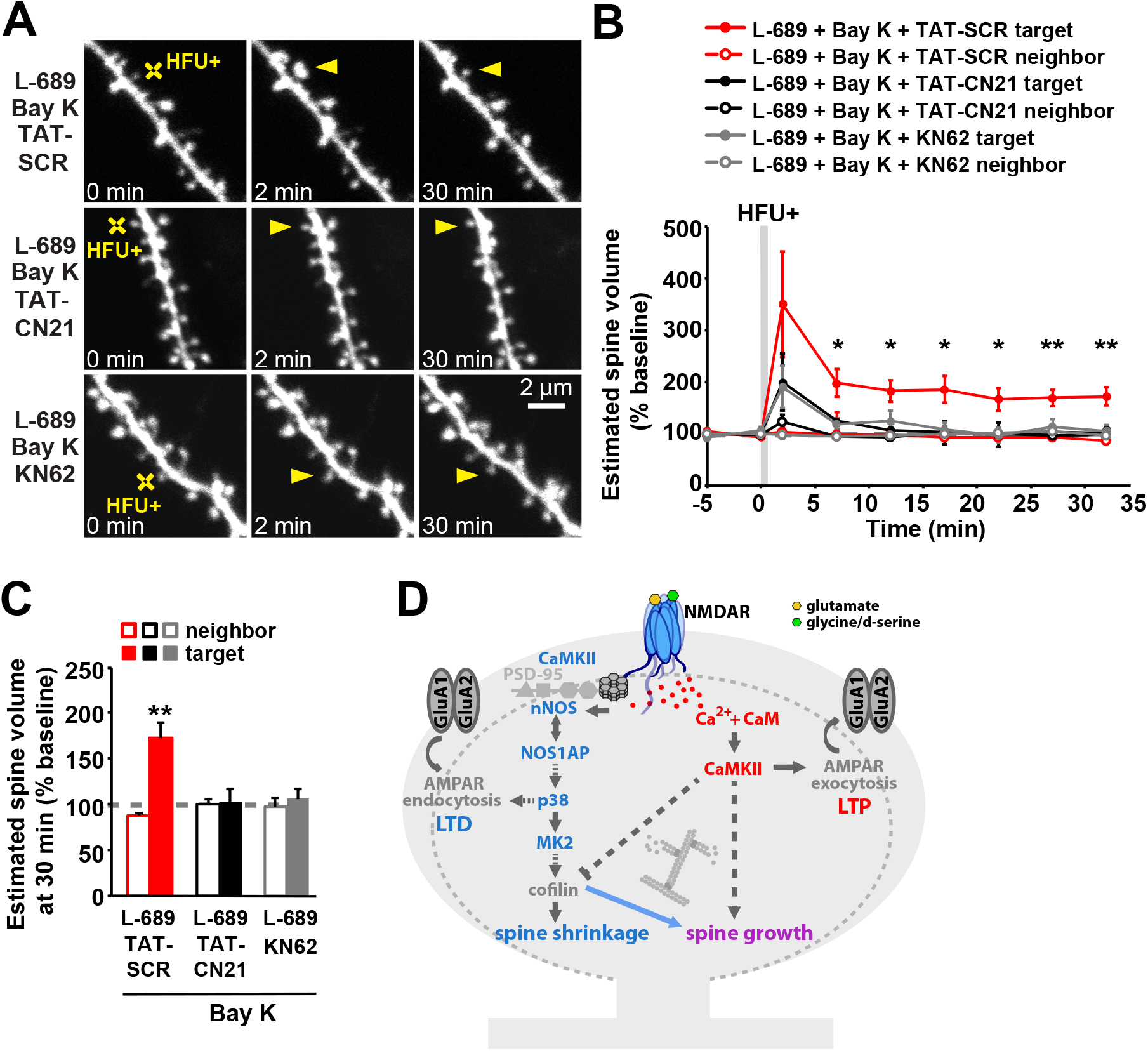
CaMKII activity is required for LTP-induced spine growth, independent of calcium source. **(A)** Images of dendrites from CA1 neurons of acute slices from P16-20 GFP-M mice before and after HFU+ stimulation (yellow cross) of individual spines (yellow arrowhead) in the presence of L-689 (10 μM) and Bay K (10 μM) or in combination with TAT-SCR (5 μM), TAT-CN21 (5 μM), or KN-62 (10 μM). **(B, C)** Spine growth in the presence of Bay K and L-689 is blocked by TAT-CN21 peptide (black filled circles/bar; 6 spines/ 6 cells) or KN-62 (gray filled circles/bars; 6 spines/6 cells), but not in the presence of control TAT-SCR peptide (red filled circles/bar; 6 spines/ 6 cells). Volume of the unstimulated neighbors (open circles/bars) was unchanged. Data are represented as mean ± SEM. *p < 0.05; **p < 0.01, ***p < 0.001. See also Figure S2. **(D)** Proposed model. In the absence of Ca^2+^-influx, plasticity-inducing stimuli activate non-ionotropic NMDAR signaling, driving dendritic spine shrinkage through cofilin dependent severing of the actin cytoskeleton. During LTP induction, severed actin filaments serve as new starting points for actin filament nucleation and branching by the Ca^2+^- and CaMKII-dependent actin modifying proteins to expand the F-actin cytoskeleton and drive spine growth.

## RESULTS

### p38 MAPK activity is required for LTP-induced spine growth, but not for LTP-induced synaptic strengthening

Glutamate binding to NMDARs initiates a signaling cascade that drives synaptic weakening (LTD) and the shrinkage of dendritic spines (sLTD), even when ion flow is blocked pharmacologically (Nabavi et al., 2013; Stein et al., 2015; Carter and Jahr, 2016; Wong and Gray, 2018). Furthermore, in the absence of ion flux, patterns of glutamatergic stimulation that would normally drive LTP and spine growth, instead drive LTD and spine shrinkage (Nabavi et al., 2013; Stein et al., 2015; Stein et al., 2020). Because the non-ionotropic NMDAR signaling pathway should also be activated with glutamate binding when ion flow through the receptor is not blocked, we wondered whether it plays a role in bidirectional synaptic plasticity. We hypothesized non-ionotropic NMDAR signaling could function as part of a built in synaptic regulatory mechanism to fine tune structural rearrangement and prevent spine overgrowth during LTP induction.

Intriguingly, inhibition of p38 MAPK activity by SB203580 did not lead to excessive HFU-induced spine growth, but instead blocked persistent spine enlargement following HFU stimulation (**Fig. 1A-C**; veh: 204 ± 13%; SB: 126 ± 12%). As p38 MAPK activation previously only had been implicated in LTD and not LTP (Zhu et al., 2002), we examined next whether expression of HFU-induced single spine LTP was normal during inhibition of p38 MAPK activity. HFU stimulation under vehicle conditions successfully induced LTP of the target spine that lasted for at least 25 min following stimulation but did not increase the uEPSC amplitude of the unstimulated neighboring spine (**Fig. 1D, F**; target: 153 ± 11%; neighbor: 93 ± 7%). Unlike spine enlargement, functional LTP induction was not blocked by SB203580 and was indistinguishable from LTP during vehicle conditions (**Fig. 1E, F**; target: 157 ± 6%; neighbor: 98 ± 9%). These results support a specific role for p38 MAPK signaling in structural, but not functional LTP.

### Non-ionotropic NMDAR signaling pathway is required for LTP-induced spine growth

The requirement of p38 MAPK for spine growth, but not for LTP, and the previously implicated role of p38 MAPK in non-ionotropic signaling during LTD and spine shrinkage led us to the unexpected hypothesis that non-ionotropic NMDAR signaling might be required for bidirectional spine structural plasticity. We have previously shown that, upstream of p38 MAPK, non-ionotropic NMDAR-dependent spine shrinkage is dependent on the interaction between NOS1AP and nNOS, and that remodeling of the actin cytoskeleton downstream of p38 MAPK is mediated by MK2 and cofilin (Stein et al., 2020). We therefore set out to test whether these molecules are also required for LTP-induced spine growth.

We tested whether NOS1AP binding to nNOS, MK2 activity, and nNOS activity, like p38 MAPK activation, are also required for persistent structural LTP. Indeed, we found that inhibition of NOS1AP binding to nNOS using L-TAT-GESV impaired persistent HFU-induced structural LTP (**Fig. 2A-C**; 126 ± 12%) compared to the inactive L-TAT-GASA control peptide (**Fig. 2A-C**; 165 ± 13%). Spine size of the unstimulated neighboring spines was not changed (**Fig. 2A-C**; L-TAT-GASA: 105 ± 4%; L-TAT-GESV: 110 ± 7%). In addition, HFU-induced spine enlargement was significantly reduced in the presence of MK2 inhibitor III compared to vehicle conditions (**Fig. 2D-F**; veh: 231 ± 27%; MK2 inhibitor III: 144 ± 20%), and no change in the size of the unstimulated neighboring spines was observed (**Fig. 2D-F**; veh: 111 ± 6%; MK2 inhibitor III: 111 ± 7%). Furthermore, in the presence of the NOS inhibitor, L-NNA, HFU-induced spine enlargement was significantly reduced compared to vehicle conditions (**Fig. 2G-I**; veh: 201 ± 28%; L-NNA: 136 ± 14%), and no change in the size of the unstimulated neighboring spines was observed (**Fig. 2G-I**; veh: 105 ± 5%; L-NNA: 110 ± 6%). Our findings strongly support a role for the non-ionotropic NMDAR signaling pathway in LTP-induced spine growth.

### Non-ionotropic NMDAR signaling in combination with VGCC-mediated Ca^2+^-influx is sufficient to drive LTP-induced spine growth

Our results support a model in which non-ionotropic NMDAR signaling leads to the p38 MAPK- and cofilin-dependent disruption of the F-actin network, which, in the absence of Ca^2+^-influx, drives spine shrinkage, but during LTP provides new actin filament nucleation and branching points for subsequent spine enlargement by Ca^2+^-dependent actin modifying and stabilizing proteins. Based on this model, we hypothesized that it should be possible to drive spine growth through a combination of non-ionotropic NMDAR signaling with NMDAR-independent Ca_2+_-influx (**Fig. 3A**).

In order to drive non-ionotropic NMDAR signaling together with voltage-gated calcium channel (VGCC)-mediated Ca^2+^-influx, the L-type Ca^2+^ channel agonist Bay K 8644 was included in the bath. Furthermore, to ensure sufficient L-type VGCC-mediated Ca^2+^-influx into the stimulated spine during glutamate uncaging, we modified our ionic conditions to increase the Ca^2+^ driving force and we increased the intensity and frequency of our high-frequency uncaging protocol (HFU+; see STAR Methods) to facilitate stronger AMPAR-dependent depolarization of the target spine. Remarkably, HFU+-stimulation in the presence of L-689 and Bay K induced long-term spine growth (**Fig. 3B-D**; L-689 + Bay K: 149 ± 10%). Importantly, adding the glutamate binding site NMDAR antagonist, CPP, which inhibits non-ionotropic NMDAR signaling, blocked HFU+-induced long-term spine growth (**Fig. 3B-D**; L-689 + Bay K + CPP: 104 ± 8%) without affecting magnitude of VGCC-mediated Ca^2+^-influx (**Fig. S1**). Size of unstimulated neighboring spines was unaffected (**Fig. 3B-D**; L-689 + Bay K: 102 ± 6%; L-689 + CPP + Bay K: 94 ± 8%), excluding acute and activity-independent effects of Bay K alone on spine morphology.

### CaMKII activity is required for long-term spine growth, independent of calcium source

CaMKII is a Ca^2+^-dependent kinase that has been extensively studied in LTP induction (Bayer and Schulman, 2019) and is required for the long-term spine growth associated with LTP (Matsuzaki et al., 2004; Murakoshi et al., 2011); **Fig. S2)**. Notably, disruption of interaction of CaMKII with the C terminus of the NMDAR interferes with LTP (Barria and Malinow, 2005; Sanhueza et al., 2011; Halt et al., 2012) and long-term spine stabilization (Hill and Zito, 2013), suggesting that the local calcium microdomains produced by calcium flow through the NMDAR may be critical in activating CaMKII during LTP-induced structural and functional plasticity.

We tested whether Ca^2+^/CaMKII-dependent signaling is also required for long-term spine growth driven by NMDAR-independent Ca^2+^ influx through VGCCs. We found that when Ca^2+^ is supplied through VGCCs and not through the NMDAR under our HFU+ stimulation conditions, long-term spine growth (**Fig. 4A, B, E**; L-689 + Bay K + TAT-SCR: 173 ± 17%) was also blocked in the presence of KN-62 (**Fig. 4A, B, E**; L-689 + Bay K + KN-62: 106 ± 11%) or TAT-CN21 (**Fig. 4A, B, E**; L689 + Bay K + TAT-CN21: 102 ± 15%), a CaMKII specific peptide inhibitor (Vest et al., 2007). Size of unstimulated neighboring spines was not affected (**Fig. 4A, B, E**; L-689 + Bay K + TAT-SCR: 87 ± 3%; L-689 + BayK + KN-62: 97 ± 10%; L-689 + Bay K + TAT-CN21: 100 ± 7%). Together, these results demonstrate the requirement of CaMKII for LTP-induced long-term spine growth, regardless of calcium source.

Finally, in order to confirm that our modified HFU+ stimulation paradigm still drives non-ionotropic NMDAR signaling induced spine shrinkage, we replaced Bay K with NBQX to block uncaging-induced AMPAR-driven spine depolarization and VGCC activation, and tested whether the same stimulation now in the absence of Ca^2+^-influx drives dendritic spine shrinkage. Indeed, HFU+ stimulation in the presence of L-689 and NBQX resulted in non-ionotropic NMDAR-dependent spine shrinkage (**Fig. 4C-E**; L-689 + NBQX: 56 ± 10%), which was inhibited if glutamate binding and thus conformational non-ionotropic NMDAR signaling was blocked with CPP (**Fig. 4C-E**; L-689 + CPP + NBQX: 100 ± 8%). Spine size of unstimulated neighbors was not changed (**Fig. 4C-E**; L-689 + NBQX: 94 ± 6%; L-689 + CPP + NBQX: 90 ± 10%). These results support our model that non-ionotropic NMDAR signaling primes the actin cytoskeleton for bidirectional structural plasticity.

## DISCUSSION

### Non-ionotropic NMDAR signaling drives bidirectional spine structural plasticity

Ion flow-independent NMDAR signaling has been implicated by many independent studies in spine shrinkage and synaptic weakening (Nabavi et al., 2013; Aow et al., 2015; Birnbaum et al., 2015; Stein et al., 2015; Carter and Jahr, 2016; Wong and Gray, 2018; Stein et al., 2020; Thomazeau et al., 2020). Here, we made the unexpected discovery that this non-ionotropic NMDAR signaling pathway is also required for spine growth during synaptic strengthening. It may appear contradictory that the same signaling pathway could support both spine shrinkage and spine growth; however, it is notable that cofilin activation, which has been implicated in spine shrinkage during LTD (Zhou et al., 2004; Hayama et al., 2013; Stein et al., 2020) is also important for long-term spine growth during LTP (Bosch et al., 2014). We propose a model for spine structural plasticity (Fig. 4F), whereby non-ionotropic NMDAR signaling leads to the p38 MAPK- and cofilin-dependent breakdown of the F-actin network, which in the absence of Ca^2+^ influx drives spine shrinkage, but with Ca^2+^-influx instead provides new actin filament nucleation and branching points for subsequent spine enlargement by Ca^2+^-dependent actin modifying and stabilizing proteins. Such a model highlights ion flux-independent NMDAR signaling as a vital component for the bidirectional structural plasticity of dendritic spines.

### Role of p38 MAPK in LTP-induced spine growth, but not synaptic strengthening

Our studies support an unexpected role for p38 MAPK, a classical LTD molecule implicated in non-ionotropic NMDAR signaling during both LTD and spine shrinkage (Nabavi et al., 2013; Stein et al., 2015), in spine growth during LTP. Notably, we found that inhibition of p38 MAPK blocked spine growth but not synaptic strengthening during LTP, at least not in the short-term. Our results are in line with previous studies reporting no role for p38 MAPK in tetanus-or pairing-induced NMDAR-dependent LTP (Zhu et al., 2002). In addition, MK2, a downstream target of p38 MAPK in the non-ionotropic NMDAR signaling pathway (Stein et al., 2020), is required for spine growth but not for LTP (Privitera et al., 2019) and NOS signaling, upstream of p38 MAPK in non-ionotropic NMDAR signaling (Stein et al., 2020), is required for both spine growth and LTP (O’Dell et al., 1994; Lu et al., 1999). Because inhibition of p38 MAPK and MK2 lead to a pharmacological dissociation of spine growth and LTP, which are normally tightly linked (Matsuzaki et al., 2004), our results suggest that these molecules are downstream of a branch point in signaling that drives synaptic strengthening and spine growth during LTP.

### Role of CaMKII in bidirectional spine structural plasticity

During LTP-induced spine growth, Ca^2+^-influx through the NMDAR leads to activation of CaMKII, which then activates small Rho GTPases, whose concerted activity drives spine enlargement via the actin regulatory proteins LIMK and Arp2/3, promoting actin polymerization and branching and spine growth (Okamoto et al., 2004; Lee et al., 2009; Murakoshi et al., 2011; Kim et al., 2013; Bosch et al., 2014; Hedrick et al., 2016; Nakahata and Yasuda, 2018; Saneyoshi et al., 2019). Our results surprisingly show that the calcium influx that drives LTP-induced spine growth does not need to enter the spine through the NMDAR, and instead can be supplied by VGCCs, as long as non-ionotropic NMDAR signaling is intact. Although it has been shown that specific local signaling microdomains are important for independently driving LTP and LTD (Zhang et al., 2018), our results suggest that local calcium microdomains produced by ion flow through the NMDAR are not critical for driving LTP-induced spine growth independent of LTD-induced spine shrinkage.

CaMKII has been extensively studied in LTP induction (Bayer and Schulman, 2019) and LTP-induced spine growth (Nishiyama and Yasuda, 2015); however, lately CaMKII also has been implicated in LTD (Coultrap et al., 2014; Woolfrey et al., 2018) and spine shrinkage driven by non-ionotropic NMDAR signaling (Stein et al., 2020). A key question is whether and how CaMKII carries out multiple roles in bidirectional spine structural plasticity. It is possible that the kinase mediates ion-flux dependent and independent signaling pathways sequentially during spine structural plasticity. Such a situation has been shown for VGCCs, in which ion flux prior to conformational signaling drives nuclear transcription (Li et al., 2016). Another possibility is that CaMKII acts simultaneously in the two pathways through different populations, perhaps one Ca^2+^/CaM bound and the other not. Indeed, synaptic activity has been proposed to activate different populations of CaMKII (Yasuda et al., 2003), which could then have different substrate specificities, as shown for Ca^2+^-bound versus autonomous CaMKII activity during LTP and LTD (Coultrap et al., 2014; Woolfrey et al., 2018).

## ACKNOWLEDGEMENTS

This work was supported by the NIH (R01 NS062736, T32 MH112507). We thank Ulli Bayer for tatCN21; Julie Culp, Jennifer Jahncke and Lorenzo Tom for support with experiments and analysis; Ulli Bayer, Juan Flores, and Samuel Petshow for critical reading of the manuscript.

## AUTHOR CONTRIBUTIONS

I.S.S., D.K.P. and K.Z. designed the experiments and wrote the manuscript. I.S.S, D.K.P., and N.C. performed the experiments and analyzed the data. All authors edited the manuscript.

---

**Figure S1 related to Figure 3:**
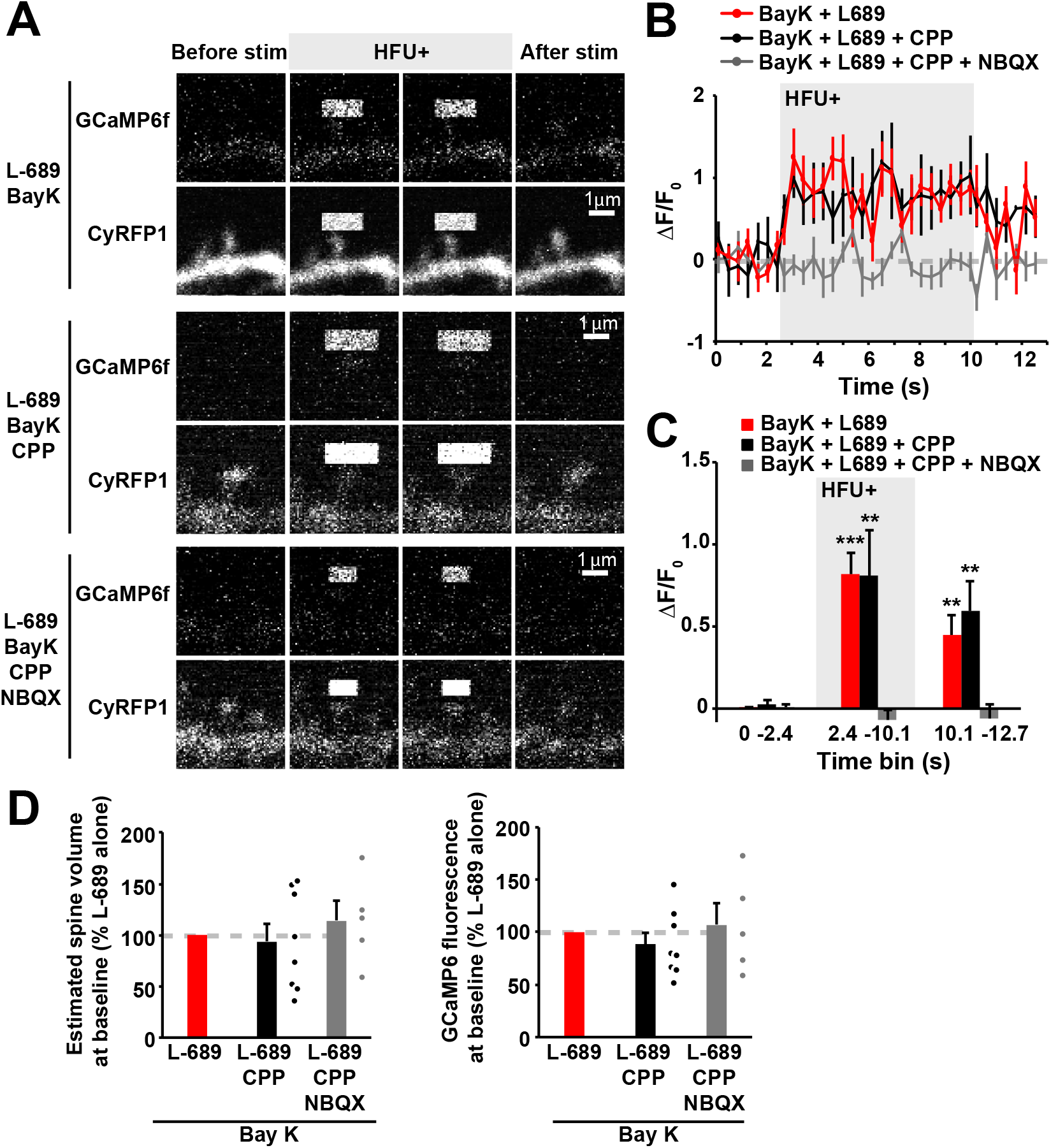
Blocking non-ionotropic NMDAR signaling with CPP does not affect the magnitude of Ca2+ influx during the HFU+ stimulation. **(A)** Images of dendrites from CA1 neurons expressing both CyRFP and GCaMP6f at DIV13-18 before, during, and after HFU+ at individual spines in the presence of L-689 (10 μM) and Bay K (10 μM), in combination with CPP (50 μM) alone, or with CPP (50 μM) and NBQX (50 μM). Middle images in each row show bleed through of uncaging laser stimulation. **(B, C)** HFU+ led to comparable levels of calcium influx into spines in the presence of L-689 and Bay K (black filled circles/bar; 11 spines/11 cells) or in combination with CPP (red filled circles/bar; 12 spines/11 cells). Calcium influx through VGCCs is blocked by inhibition of AMPARs with NBQX (gray filled circles/bar; 10 spines/9 cells). (D) Left: No difference in baseline volume of spines in Bay K, L-689, and CPP (red; 8 spines/8 cells); and Bay K, L-689, CPP, and NBQX (gray; 5 spines/5 cells) relative to those exposed to Bay K and L-689 alone (black; 8 spines/8 cells). Right: No difference in baseline GCaMP6 fluorescence of spines in Bay K, L-689, and CPP (red; 8 spines/ 8 cells); and Bay K, L-689, CPP, and NBQX (gray; 5 spines/5 cells) relative to that for Bay K and L-689 alone (black; 8 spines/8 cells). Data are represented as mean +/- SEM. *p < 0.05; **p < 0.01, ***p < 0.001.

**Figure S2, related to Figure 4:**
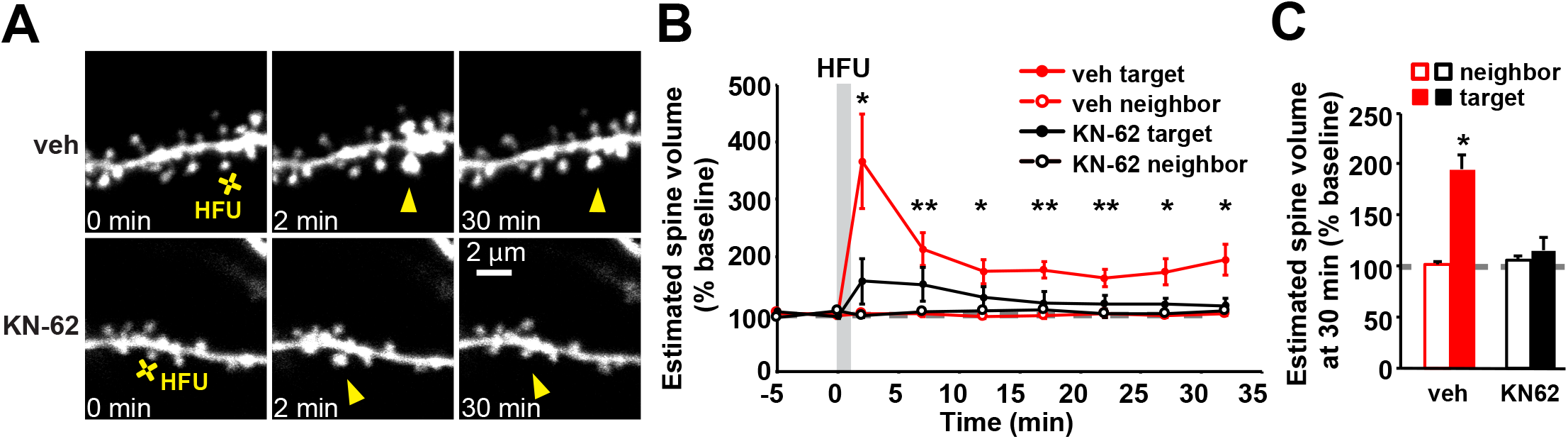
Inhibition of CaMKII blocks LTP-induced long-term spine growth. **(A)** Images of dendrites from CA1 neurons of P16-20 acute slices from GFP-M mice before and after HFU stimulation (yellow cross) of individual spines during vehicle conditions and in the presence of CaMKII inhibitor KN-62 (10 μM). **(B, C)** HFU-induced dendritic spine growth (vehicle, red filled circles/bar; 8 spines/8 cells) was prevented by KN-62 (black filled circles/bar; 6 spines/6 cells). Volume of the unstimulated neighbors did not change (open bars). Data are represented as mean +/- SEM. *p < 0.05; **p < 0.01, ***p < 0.001.

## REFERENCES

Aow J, Dore K, Malinow R (2015) Conformational signaling required for synaptic plasticity by the NMDA receptor complex. Proc Natl Acad Sci U S A 112:14711–14716.

Barria A, Malinow R (2005) NMDA receptor subunit composition controls synaptic plasticity by regulating binding to CaMKII. Neuron 48:289–301.

Bayer KU, Schulman H (2019) CaM Kinase: Still Inspiring at 40. Neuron 103:380–394.

Birnbaum JH, Bali J, Rajendran L, Nitsch RM, Tackenberg C (2015) Calcium flux-independent NMDA receptor activity is required for Abeta oligomer-induced synaptic loss. Cell Death Dis 6:e1791.

Bosch M, Castro J, Saneyoshi T, Matsuno H, Sur M, Hayashi Y (2014) Structural and molecular remodeling of dendritic spine substructures during long-term potentiation. Neuron 82:444–459.

Carter BC, Jahr CE (2016) Postsynaptic, not presynaptic NMDA receptors are required for spike-timing-dependent LTD induction. Nat Neurosci 19:1218–1224.

Chen TW, Wardill TJ, Sun Y, Pulver SR, Renninger SL, Baohan A, Schreiter ER, Kerr RA, Orger MB, Jayaraman V, Looger LL, Svoboda K, Kim DS (2013) Ultrasensitive fluorescent proteins for imaging neuronal activity. Nature 499:295–300.

Coultrap SJ, Freund RK, O’Leary H, Sanderson JL, Roche KW, Dell’Acqua ML, Bayer KU (2014) Autonomous CaMKII mediates both LTP and LTD using a mechanism for differential substrate site selection. Cell Rep 6:431–437.

Dore K, Aow J, Malinow R (2015) Agonist binding to the NMDA receptor drives movement of its cytoplasmic domain without ion flow. Proc Natl Acad Sci U S A 112:14705–14710.

Feng G, Mellor RH, Bernstein M, Keller-Peck C, Nguyen QT, Wallace M, Nerbonne JM, Lichtman JW, Sanes JR (2000) Imaging neuronal subsets in transgenic mice expressing multiple spectral variants of GFP. Neuron 28:41–51.

Ferreira JS, Papouin T, Ladepeche L, Yao A, Langlais VC, Bouchet D, Dulong J, Mothet JP, Sacchi S, Pollegioni L, Paoletti P, Oliet SHR, Groc L (2017) Co-agonists differentially tune GluN2B-NMDA receptor trafficking at hippocampal synapses. Elife 6.

Halt AR, Dallapiazza RF, Zhou Y, Stein IS, Qian H, Juntti S, Wojcik S, Brose N, Silva AJ, Hell JW (2012) CaMKII binding to GluN2B is critical during memory consolidation. Embo j 31:1203–1216.

Hayama T, Noguchi J, Watanabe S, Takahashi N, Hayashi-Takagi A, Ellis-Davies GC, Matsuzaki M, Kasai H (2013) GABA promotes the competitive selection of dendritic spines by controlling local Ca2+ signaling. Nat Neurosci 16:1409–1416.

Hayashi-Takagi A, Yagishita S, Nakamura M, Shirai F, Wu YI, Loshbaugh AL, Kuhlman B, Hahn KM, Kasai H (2015) Labelling and optical erasure of synaptic memory traces in the motor cortex. Nature 525:333–338.

Hedrick NG, Harward SC, Hall CE, Murakoshi H, McNamara JO, Yasuda R (2016) Rho GTPase complementation underlies BDNF-dependent homo- and heterosynaptic plasticity. Nature 538:104–108.

Hill TC, Zito K (2013) LTP-induced long-term stabilization of individual nascent dendritic spines. J Neurosci 33:678–686.

Holtmaat AJ, Trachtenberg JT, Wilbrecht L, Shepherd GM, Zhang X, Knott GW, Svoboda K (2005) Transient and persistent dendritic spines in the neocortex in vivo. Neuron 45:279–291.

Kim IH, Racz B, Wang H, Burianek L, Weinberg R, Yasuda R, Wetsel WC, Soderling SH (2013) Disruption of Arp2/3 results in asymmetric structural plasticity of dendritic spines and progressive synaptic and behavioral abnormalities. J Neurosci 33:6081–6092.

Lai CSW, Adler A, Gan WB (2018) Fear extinction reverses dendritic spine formation induced by fear conditioning in the mouse auditory cortex. Proc Natl Acad Sci U S A 115:9306–9311.

Laviv T, Kim BB, Chu J, Lam AJ, Lin MZ, Yasuda R (2016) Simultaneous dual-color fluorescence lifetime imaging with novel red-shifted fluorescent proteins. Nat Methods 13:989–992.

Lee SJ, Escobedo-Lozoya Y, Szatmari EM, Yasuda R (2009) Activation of CaMKII in single dendritic spines during long-term potentiation. Nature 458:299–304.

Li B, Tadross MR, Tsien RW (2016) Sequential ionic and conformational signaling by calcium channels drives neuronal gene expression. Science 351:863–867.

Lu YF, Kandel ER, Hawkins RD (1999) Nitric oxide signaling contributes to late-phase LTP and CREB phosphorylation in the hippocampus. J Neurosci 19:10250–10261.

Matsuzaki M, Honkura N, Ellis-Davies GC, Kasai H (2004) Structural basis of long-term potentiation in single dendritic spines. Nature 429:761–766.

Murakoshi H, Wang H, Yasuda R (2011) Local, persistent activation of Rho GTPases during plasticity of single dendritic spines. Nature 472:100–104.

Nabavi S, Kessels HW, Alfonso S, Aow J, Fox R, Malinow R (2013) Metabotropic NMDA receptor function is required for NMDA receptor-dependent long-term depression. Proc Natl Acad Sci U S A 110:4027–4032.

Nakahata Y, Yasuda R (2018) Plasticity of Spine Structure: Local Signaling, Translation and Cytoskeletal Reorganization. Front Synaptic Neurosci 10:29.

Nishiyama J, Yasuda R (2015) Biochemical Computation for Spine Structural Plasticity. Neuron 87:63–75.

O’Dell TJ, Huang PL, Dawson TM, Dinerman JL, Snyder SH, Kandel ER, Fishman MC (1994) Endothelial NOS and the blockade of LTP by NOS inhibitors in mice lacking neuronal NOS. Science 265:542–546.

Okamoto K, Nagai T, Miyawaki A, Hayashi Y (2004) Rapid and persistent modulation of actin dynamics regulates postsynaptic reorganization underlying bidirectional plasticity. Nat Neurosci 7:1104–1112.

Pologruto TA, Sabatini BL, Svoboda K (2003) ScanImage: flexible software for operating laser scanning microscopes. Biomed Eng Online 2:13.

Privitera L, Hogg EL, Gaestel M, Wall MJ, Corrêa SAL (2019) The MK2 cascade regulates mGluR-dependent synaptic plasticity and reversal learning. Neuropharmacology 155:121–130.

Saneyoshi T, Matsuno H, Suzuki A, Murakoshi H, Hedrick NG, Agnello E, O’Connell R, Stratton MM, Yasuda R, Hayashi Y (2019) Reciprocal Activation within a Kinase-Effector Complex Underlying Persistence of Structural LTP. Neuron 102:1199–1210 e1196.

Sanhueza M, Fernandez-Villalobos G, Stein IS, Kasumova G, Zhang P, Bayer KU, Otmakhov N, Hell JW, Lisman J (2011) Role of the CaMKII/NMDA receptor complex in the maintenance of synaptic strength. J Neurosci 31:9170–9178.

Stein IS, Zito K (2019) Dendritic Spine Elimination: Molecular Mechanisms and Implications. Neuroscientist 25:27–47.

Stein IS, Gray JA, Zito K (2015) Non-Ionotropic NMDA Receptor Signaling Drives Activity-Induced Dendritic Spine Shrinkage. J Neurosci 35:12303–12308.

Stein IS, Park DK, Flores JC, Jahncke JN, Zito K (2020) Molecular Mechanisms of Non-ionotropic NMDA Receptor Signaling in Dendritic Spine Shrinkage. The Journal of Neuroscience 40:3741–3750.

Stoppini L, Buchs PA, Muller D (1991) A simple method for organotypic cultures of nervous tissue. J Neurosci Methods 37:173–182.

Thomazeau A, Bosch M, Essayan-Perez S, Barnes SA, De Jesus-Cortes H, Bear MF (2020) Dissociation of functional and structural plasticity of dendritic spines during NMDAR and mGluR-dependent long-term synaptic depression in wild-type and fragile X model mice. Molecular Psychiatry.

Vest RS, Davies KD, O’Leary H, Port JD, Bayer KU (2007) Dual mechanism of a natural CaMKII inhibitor. Mol Biol Cell 18:5024–5033.

Wong JM, Gray JA (2018) Long-Term Depression Is Independent of GluN2 Subunit Composition. J Neurosci 38:4462–4470.

Woods G, Zito K (2008) Preparation of gene gun bullets and biolistic transfection of neurons in slice culture. J Vis Exp.

Woods GF, Oh WC, Boudewyn LC, Mikula SK, Zito K (2011) Loss of PSD-95 enrichment is not a prerequisite for spine retraction. J Neurosci 31:12129–12138.

Woolfrey KM, O’Leary H, Goodell DJ, Robertson HR, Horne EA, Coultrap SJ, Dell’Acqua ML, Bayer KU (2018) CaMKII regulates the depalmitoylation and synaptic removal of the scaffold protein AKAP79/150 to mediate structural long-term depression. J Biol Chem 293:1551–1567.

Yasuda R, Sabatini BL, Svoboda K (2003) Plasticity of calcium channels in dendritic spines. Nat Neurosci 6:948–955.

Zhang L, Zhang P, Wang G, Zhang H, Zhang Y, Yu Y, Zhang M, Xiao J, Crespo P, Hell JW, Lin L, Huganir RL, Zhu JJ (2018) Ras and Rap Signal Bidirectional Synaptic Plasticity via Distinct Subcellular Microdomains. Neuron 98:783–800 e784.

Zhou Q, Homma KJ, Poo MM (2004) Shrinkage of dendritic spines associated with long-term depression of hippocampal synapses. Neuron 44:749–757.

Zhu JJ, Qin Y, Zhao M, Van Aelst L, Malinow R (2002) Ras and Rap control AMPA receptor trafficking during synaptic plasticity. Cell 110:443–455.

